# Dual spatially resolved transcriptomics for SARS-CoV-2 host-pathogen colocalization studies in humans

**DOI:** 10.1101/2022.03.14.484288

**Authors:** Hailey Sounart, Enikő Lázár, Yuvarani Masarapu, Jian Wu, Tibor Várkonyi, Tibor Glasz, András Kiss, Erik Borgström, Andrew Hill, Aleksandra Jurek, Anezka Niesnerová, Henrik Druid, Olaf Bergmann, Stefania Giacomello

**Author notes:** First shared authors (Hailey Sounart, Enikő Lázár). Senior shared authors (Olaf Bergmann, Stefania Giacomello).

## Abstract

To advance our understanding of cellular host-pathogen interactions, technologies that facilitate the co-capture of both host and pathogen spatial transcriptome information are needed. Here, we present an approach to simultaneously capture host and pathogen spatial gene expression information from the same formalin-fixed paraffin embedded (FFPE) tissue section using the spatial transcriptomics technology. We applied the method to COVID-19 patient lung samples and enabled the dual detection of human and SARS-CoV-2 transcriptomes at 55 μm resolution. We validated our spatial detection of SARS-CoV-2 and identified an average specificity of 94.92% in comparison to RNAScope and 82.20% in comparison to *in situ* sequencing (ISS). COVID-19 tissues showed an upregulation of host immune response, such as increased expression of inflammatory cytokines, lymphocyte and fibroblast markers. Our colocalization analysis revealed that SARS-CoV-2^+^ spots presented shifts in host RNA metabolism, autophagy, NFκB, and interferon response pathways. Future applications of our approach will enable new insights into host response to pathogen infection through the simultaneous, unbiased detection of two transcriptomes.

## Introduction

Much is still unknown about how hosts react to pathogens and how pathogen infection underlies various biological processes and disease states. Although single-cell transcriptomics methods have improved the elucidation of cell-type specific effects caused by pathogens and how these relate to disease outcomes^1,2^, such approaches remove pathogens and host cells from their natural environment, limiting the study of complex dynamics of localized infections. To gain insights on the localized host response to pathogen infection, technologies that allow the co-capture of both host and pathogen spatial transcriptome information are needed. Moreover, there is often the need to work with formalin-fixed paraffin embedded (FFPE) tissue blocks to neutralize the pathogen and, when studying human infectious diseases, to access biobanks where samples are deposited. For instance, in light of the COVID-19 pandemic, human and SARS-CoV-2 transcriptome information was spatially captured from FFPE lung tissues to study human host response to SARS-CoV-2 viral infection in the lung^3–9^, providing new insights into the heterogeneous viral distribution and host response to infection. However, although there are currently some FFPE compatible spatially-resolved transcriptomics methods available^10–18^, such methods are limited by either providing only a partial view of the full transcriptome^11–14^ or having low tissue area throughput, due to either long experimental times^10–15,16^ or having to rely on the selection of predefined tissue regions of interest^19^.

Here, we present a spatially-resolved transcriptomics strategy to unbiasedly explore the transcriptome-wide landscape of two transcriptomes using FFPE tissues. We leveraged the high-throughput, sequencing-based commercially available spatial transcriptomic (ST) platform^20^ and introduced the co-detection of a second transcriptome, that of the SARS-CoV-2 virus, to the human one. We tested the potential of such an approach through the dual capture of human and SARS-CoV-2 viral transcriptomes at 55mμm (~1-10 cells) spatial resolution in COVID-19 patient lung FFPE tissues. We validated our spatial detection of SARS-CoV-2 with targeted transcriptome technologies RNAScope^11^ and *in situ* sequencing (ISS)^14,21,22^. With our approach, we identified both general immune response signatures and tissue-specific processes evoked by the virus, such as the domination of plasma cells, activated fibroblasts, and inflammatory cytokines in COVID-19 lung tissues due to prolonged SARS-CoV-2 infection. A prominent feature of our method is the colocalization of human and viral gene expression information that allows an understanding of human tissue response to viral infection by comparing areas with and without the presence of viral RNA in the same tissue section. Such an approach uncovered several genes involved in RNA metabolism, autophagy, NFκB, and interferon-response pathways to be differentially expressed in viral active areas, potentially shedding new light on COVID-19 pathogenesis. Our strategy opens up the possibility of spatially studying host response to pathogen infections through the simultaneous, unbiased detection of two transcriptomes.

## Results

### Spatial Transcriptomics enables the simultaneous capture of human and SARS-CoV-2 spatial transcriptomes

To study the localized infection effects caused by SARS-CoV-2 in human lungs, we advanced the Visium Spatial Gene Expression assay for FFPE tissues^20^ to simultaneously capture human and SARS-CoV-2 whole transcriptome (WT) information at a 55 μm resolution. Specifically, we analyzed 16,688 human genes and 10 SARS-CoV-2 gene transcripts in total across 13 tissue sections from 5 patient lung tissue samples, 3 from COVID-19 patients (i.e., 1C, 2C, 3C) and 2 from control patients (i.e., 4nC, 5nC) (Figure 1a, Supplementary Figure 1, Supplementary Table 1, Supplementary Table 2). First, we verified the specificity of the SARS-CoV-2 probes (S) for capturing SARS-CoV-2 transcripts only. To do so, we applied both human WT probes (H) and spike-in SARS-CoV-2 probes to control tissue sections. We did not identify any SARS-CoV-2 transcripts above background levels (see Methods), demonstrating the SARS-CoV-2 probes were specific to capture SARS-CoV-2 information (Figure 1b, Supplementary Figure 2).

**Figure 1.**
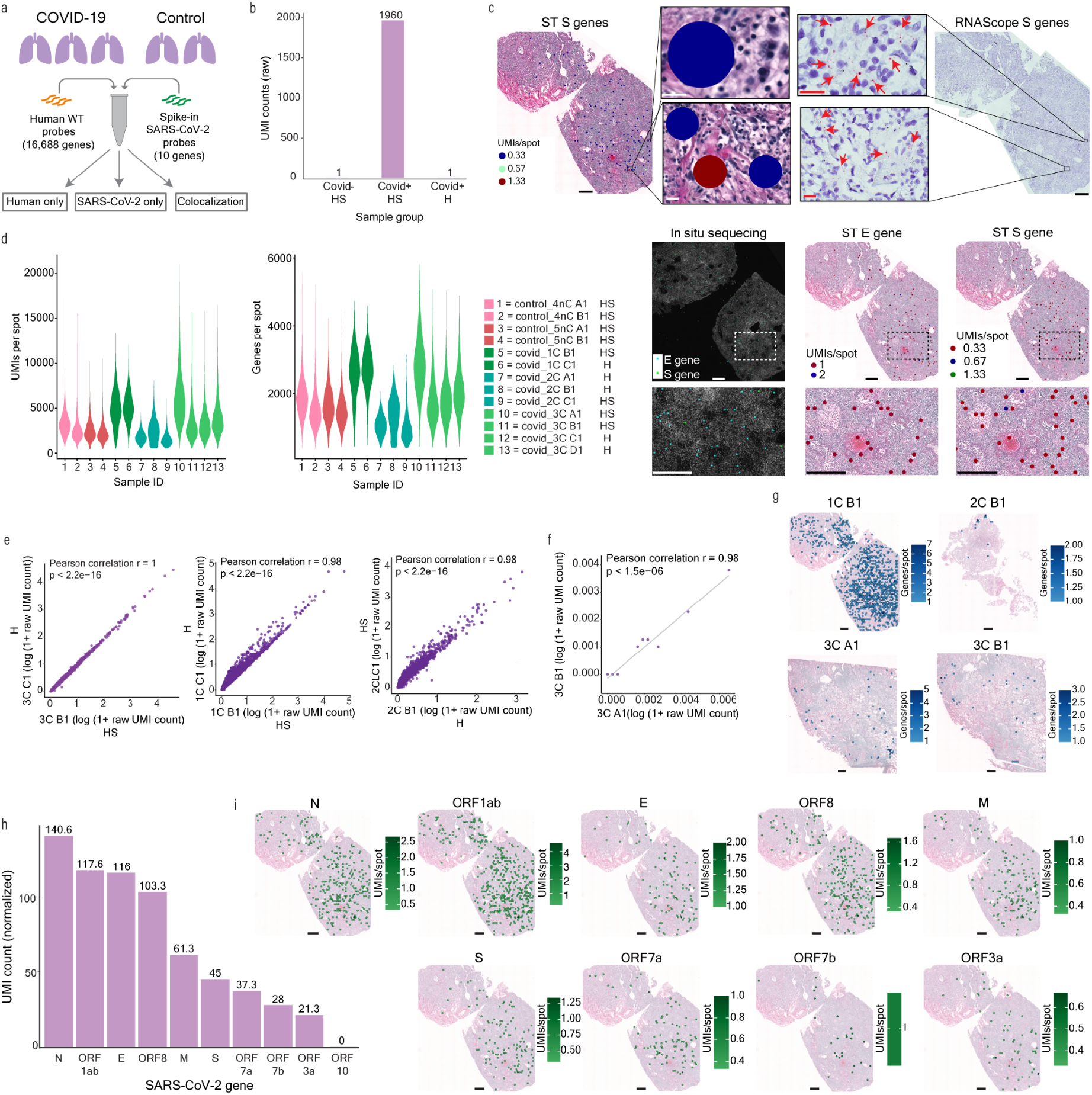
Dual detection of human and SARS-CoV-2 transcriptomes by Spatial Transcriptomics. (a) Overview of the study. (b) SARS-CoV-2 detection in control sections with human and SARS-CoV-2 probes added (HS), in COVID-19 sections with human and SARS-CoV-2 probes added (HS), and in COVID-19 sections with only human probes added (H). (c) Distribution of UMI and gene counts per capture spot across patient sample sections. (d) Pearson correlation of average human gene expression between consecutive sections for each sample, one section with human and SARS-CoV-2 probes added (HS) and the other with only human probes added (H). P-value < 2.2e-16. (e) SARS-CoV-2 S gene detection by ST and RNAScope in a consecutive section. SARS-CoV-2 S & E genes detection by ST and *in situ* sequencing (ISS). ST and RNAScope full tissue section scale bars are 550μm. RNAScope and ST S gene zoomed in panel scale bars are 20μm. All ISS and ST panel scale bars are 550μm. (f) Average SARS-CoV-2 transcriptome capture between consecutive sections. (g) SARS-CoV-2 genes per capture spot across each COVID-19 sample. Scale bars are 500μm. (h) Abundance (total normalized UMI counts) of the SARS-CoV-2 genes across all COVID-19 samples. (i) The spatial distribution of UMI counts per capture spot of each SARS-CoV-2 gene for COVID-19 sample 1C. Scale bars are 500μm.

To independently validate the viral detection by our set of SARS-CoV-2 probes, we compared the ST viral detection to the viral detection offered by the orthogonal imaging-based RNAScope technology^11^ using consecutive sections. Specifically, we compared the S gene signal detected by ST and RNAScope across all COVID-19 and control samples. To systemically and unbiasedly analyze all our samples, we developed an automated signal detection computational pipeline across all platforms (see Methods, Supplementary Figures 3-5). Using our computational approach, we found an average specificity of the ST method of 94.92% (1C: 86.86%, 2C: 99.37%, 3C: 98.53%) (Figure 1c). Furthermore, we performed a second validation of the ST S gene detection, and a validation of the ST E gene, by using *in situ* sequencing (ISS)^14,21,22^ in the sample with the highest viral load, 1C, using the same automated computational pipeline (see Methods). Despite several sections in between (~300 μm) the ST and ISS sections, we observed an overall similar viral distribution trend between ST and ISS sections and an average specificity of 82.20% (83.65% for the E gene and 80.74% for the S gene) (Figure 1c, Supplementary Figure 6), confirming that the ST method can capture SARS-CoV-2 information accurately.

Subsequently, we wanted to understand if the addition of the SARS-CoV-2 probes impacted the quality of the human gene expression information captured. To this end, we analyzed consecutive COVID-19 sections with human WT probes and spike-in SARS-CoV-2 probes versus only human WT probes. Across COVID-19 and control tissue sections, we generated a dataset consisting of a total of 37,754 spots, with an average of ~2,013 unique human genes and ~3,809 unique human molecules per spot, respectively (Figure 1d). We captured very similar human gene expression profiles between sample replicate sections and across most samples, both with and without SARS-CoV-2 probes added (*r*= 0.98-1, p-value < 2.2e-16) (Figure 1d-e, Supplementary Figure 7). Overall, these results demonstrate highly reproducible capture of human transcriptomic information and the specificity of the SARS-CoV-2 probes in detecting the SARS-CoV-2 transcriptome without interfering with the capture efficiency of the human transcripts.

### SARS-CoV-2 genes have different expression levels but follow a similar spatial distribution

In COVID-19 sections, 9.5% of spots (i.e., 1,132 spots in total) presented SARS-CoV-2 transcriptional signal with highly reproducible capture of SARS-CoV-2 gene expression between consecutive sections (*r*=0.98, p-value < 1.5e-6, Figure 1f). Overall, we captured up to 9 different SARS-CoV-2 genes (Supplementary Table 3) with an average per spot of ~1.7 unique molecules and ~ 1.5 unique genes, respectively. These relative low levels of viral load are likely associated with the longer disease duration (13-17 days) of these patients (Supplementary Table 2) in agreement with several studies that observed lower, or even undetectable, viral load in COVID-19 patients with longer survival times^3–5,7,23^. By looking at the overall distribution of SARS-CoV-2^+^ spots in COVID-19 samples, we observed a wide range of percent SARS-CoV-2^+^ spots in a COVID-19 sample section: 33.6% for 1C, 1.1% for 2C, and 1.0-1.6% for 3C (Figure 1g, Supplementary Table 3). Others have observed such inter-sample viral load heterogeneity^3,4,7,9^ and even heterogeneous distribution within the same tissue sample^7^. In this regard, we also observed varied abundances of the different SARS-CoV-2 gene transcripts across all three COVID-19 samples (Figure 1h, Supplementary Figure 8). For example, across our COVID-19 sample sections, N was the highest expressed SARS-CoV-2 gene while ORF10 was not detected at all, in line with previous reports of N as the most abundant subgenomic RNA (sgRNA)^24,25^ and ORF10 as consistently either absent or the lowest sgRNA detected^24,25^. The abundance trend of the remaining SARS-CoV-2 genes (M, E, S, ORF1ab, ORF3a, ORF7a, ORF7b, ORF8) varied across the four COVID-19 sample sections (Supplementary Figure 8) and factors driving these differences remain to be further investigated in the literature. Although the SARS-CoV-2 transcripts differed in their abundances across genes, we observed a fairly even spatial distribution of each gene across samples 1C and 3C, while for 2C the transcripts showed a more localized spatial distribution (Figure 1i, Supplementary Figures 9-10). Variation in the SARS-CoV-2 gene abundances could be influenced by the SARS-CoV-2 probes binding to both viral genomic RNA (gRNA) and subgenomic RNA (sgRNA), as previous studies observed sgRNA abundance variation^24,25^ and such differences could be reflected in our SARS-CoV-2 transcriptomic data.

### COVID-19 induces lymphocyte infiltration and expressional shifts in lung myeloid, fibroblast, and alveolar epithelial cells

To explore the human lung cellular landscape in response to SARS-CoV-2 infection, we performed unsupervised, joint graph-based clustering of spatial transcriptome data collected from both COVID-19 and control sections and found six distinct clusters (Figure 2a). Investigation into the differentially expressed (DE) genes of the six clusters revealed all of them to be a mixture of different cell types, however, we identified clear DE gene signatures for clusters dominated by myeloid cells (cluster 1), endothelial cells (cluster 2), B-cells/plasma cells (cluster 3), epithelial cells (cluster 4), and fibroblasts (cluster 6) (Figure 2b-c, Supplementary Figure 11, Supplementary Table 4). Cluster 5 was characterized by DE genes specific for endothelial, fibroblast and smooth muscle cells, where further sub clustering of this group resulted in three subclusters (subcluster 1: fibroblast-dominated, subcluster 2: smooth muscle cell-dominated, subcluster 3: mixture of endothelial and immune cells), in line with our previous observations (Supplementary Table 5). SARS-CoV-2^+^ spots appeared throughout different morphological areas without obvious enrichment in any of the spatial clusters (Figure 2a), possibly explained by the resolution of spatial transcriptomics not yet reaching the single cell level, and in agreement with the uniform spatial gene expression of the SARS-CoV-2 genes across the same tissue section (Figure 1i, Supplementary Figure 10)

**Figure 2.**
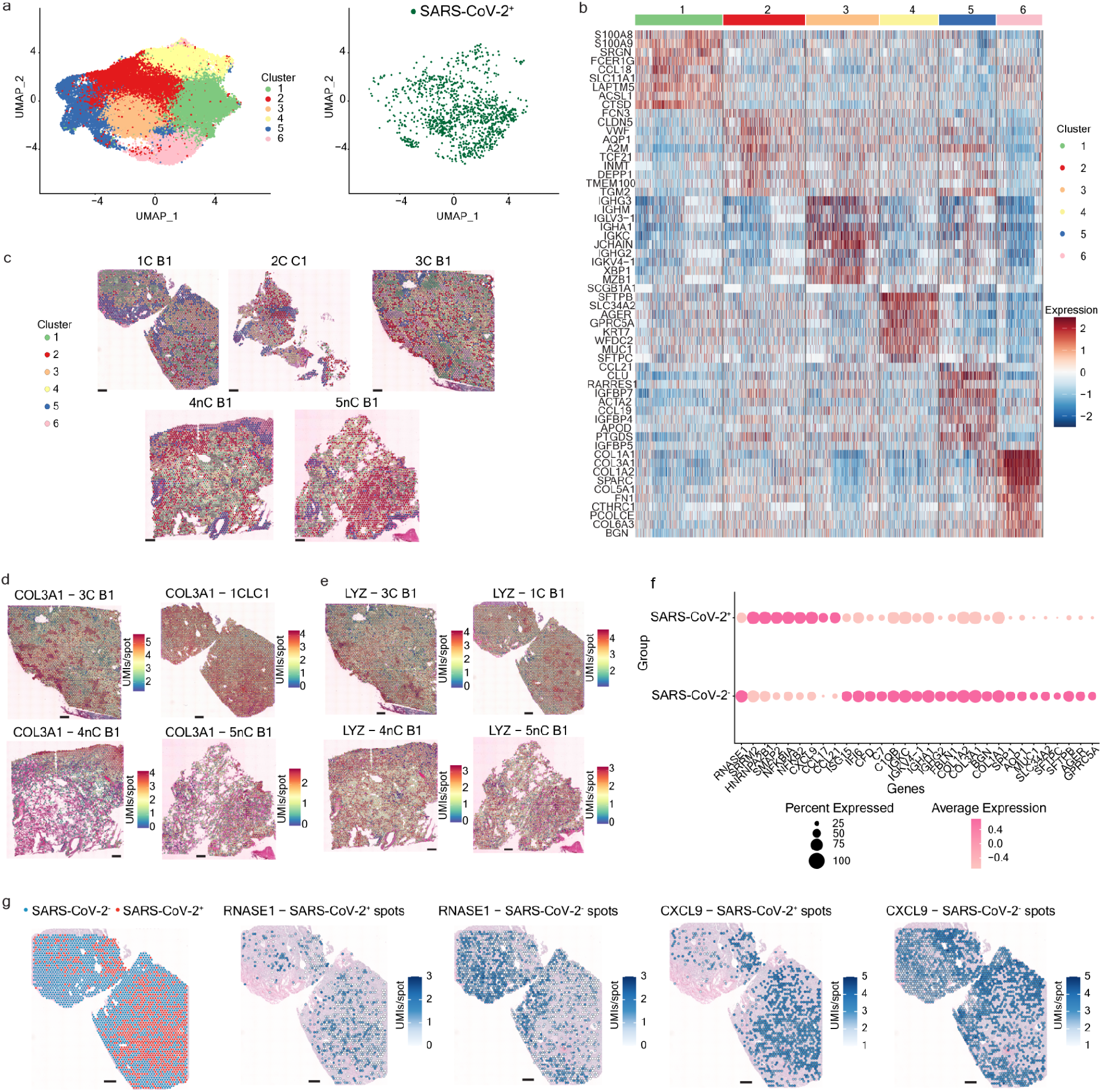
Human host response to SARS-CoV-2 infection. (a) Clustering of the human transcriptome data across COVID-19 and control sections reveals 6 distinct clusters with SARS-CoV-2^+^ spots distributed throughout the clusters. (b) Differential genes per cluster across COVID-19 and control sections. (c) Spatial distribution of the clusters on COVID-19 and control sections. Scale bars are 500μm. (d-e) Spatial distribution of genes upregulated in COVID-19 sections, COL3A1 (d) and LYZ (e). Scale bars are 500μm. (f) Dotplot depicting differential expression of human genes in SARS-CoV-2^+^ and SARS-CoV-2^-^ spots in COVID-19 sections. (g) Spatial distribution of RNase1, downregulated in SARS-CoV-2^+^ spots, and CXCL9, upregulated in SARS-CoV-2^+^ spots, in COVID-19 sample 1C. Scale bars are 500μm.

All three COVID-19 lung samples included in this study represented the late-phase pneumonia stage (between 13-17 days post-infection) and showed consistent histopathological features with organizing diffuse alveolar damage, extensive fibrosis, and leukocyte infiltration, accompanied by low viral load^4,7,23^ (Supplementary Figure 1). We found that these substantial structural differences between the COVID-19 and control lung sections were also indicated in the transcriptome data. Specifically, DE genes for COVID-19 lung sections were dominated by signatures of plasma cells (IGHG3, IGKC, IGHM, JCHAIN, IGHG2, IGKV4-1, IGLV3-1, IGHA1), activated fibroblasts (COL1A1, COL1A2, COL3A1), inflammatory cytokines (CXCL9, CCL18) and complement factors (C1QB, C1QC), reflecting the overall tissue response to a prolonged SARS-CoV-2 infection (Figure 2d, Supplementary Table 6).

We did not find viral entry factors ACE2, TMPRSS2, PCK5, or PCSK7 to be differentially expressed between COVID-19 and control lung sections, in line with a recent study^3^. However, we found three other known SARS-CoV-2 entry factors, CTSL^26,27^, CTSB^26^, and NRP1^28,29^ to be upregulated in the COVID-19 lungs (Supplementary Table 6). Previous studies^30,31^ observed that CTSL and CTSB have increased expression in COVID-19 lungs, with SARS-CoV-2 infection demonstrated to promote CTSL expression that in turn enhances viral infection^27^.

Within the myeloid-cell rich cluster 1 of the COVID-19 tissue samples, we observed selective up- (CD163, F13A1, CD14, LYZ, APOE, C1QA, B2M) or downregulation (PPARG, VCAN, FCN1, YAP1, FCGR3A) of certain marker genes, previously used to annotate distinct monocyte-macrophage lineage subsets in single cell transcriptomic datasets of COVID-19 lungs^3^, implicating a shift in myeloid cell subtypes during disease progression (Supplementary Table 7). Besides many consensus fibrosis markers (COL1A1, COL3A1, COL5A1, SPP1, FN1, POSTN), we found the CTHRC1 gene, recently described as a marker of pathological, pulmonary fibrosis-associated fibroblasts^32^, to be highly expressed in the fibroblast-rich cluster 6 of COVID-19 lungs (Supplementary Table 7). At the same time, we identified markers of alveolar fibroblasts, thought to be a cellular source of the CTHRC1^+^ fibroblast subpopulation^32^, as either mildly downregulated (TCF21, PDGFRA) or upregulated (NPNT1) in the same cluster (Supplementary Table 7).

To investigate the biological changes occurring in the epithelial cell compartments, we performed further sub-clustering of cluster 4 (Supplementary Table 8). In the pneumocyte-dominated subcluster 1, COVID-19 lung tissue showed markedly (SFTPC, SLC34A2, MUC1, LYZ) or moderately (LAMP3, PGC, NAPSA, CEBPA, LPCAT1, SDC1, NKX2-1, PIGR, ABCA3, ALPL) elevated expression of type 2 alveolar epithelial (AT2) cell markers, while the type 1 alveolar epithelial (AT1) cell markers appeared to be either slightly down- (KRT7, AGER, FSTL3, SCNN1G) or upregulated (CYP4B1, ICAM1, AQP4, RTKN2, EMP2, GPRC5A, AQP3, CAV1) (Figure 2e, Supplementary Table 9). These changes might point at a hyperplastic expansion of AT2 cells in the diseased tissue, consistent with microscopic observations made in lungs with longer disease duration^33,34^ and characterized by low viral load^4^.

### Colocalization analysis of SARS-CoV-2 and human transcriptomes identifies dysregulation of RNA metabolism and NFκB pathway activation

With the possibility of separately capturing the human and SARS-CoV-2 spatial transcriptomes, our approach allowed us to conduct colocalization studies and identify host gene expression changes caused by the active presence of the viral mRNA in lung cells at 55 μm resolution. By comparing the human gene expression patterns in SARS-CoV-2^+^ and SARS-CoV-2^-^ spots only in COVID-19 tissue sections (Supplementary Table 10), we revealed several genes involved in RNA metabolism to be affected by the presence of the viral mRNA (Figure 2f, Supplementary Table 10). We found SRRM2, a component of the spliceosome, to be upregulated in SARS-CoV-2^+^ spots (Figure 2f, Supplementary Table 10). SRRM2 plays a central role in nuclear speckle formation and thus in the replication, splicing and trafficking of Herpes Simplex Virus and Influenza A virus^35^, which suggests a potentially similar role in the processing of the SARS-CoV-2 virus. A study described HNRNPA2/B1, another upregulated gene involved in the packaging of nascent pre-mRNA, in the formation of cytoplasmic stress granules responsible for the assembly of the nucleocapsid protein and genomic RNA of SARS-CoV-2^36^ (Supplementary Table 10). Another study showed that the NSP1 of SARS-CoV-2 facilitates viral RNA processing and blocks effective IFNβ expression through directly binding HNRNPA2/B1 and redistributing it between the nucleus and cytoplasm^37^. Direct modulation of SMMR2 and HNRNPA2/B1 expression levels in the host cells might be an alternative mechanism for the SARS-CoV-2 virus to ensure proper synthesis, assembly, and further spreading of viral particles. Notably, we found the RNase1 gene to be downregulated in SARS-CoV-2^+^ spots, potentially blocking degradation of viral RNA in the environment of actively infected cells (Figure 2f-g, Supplementary Table 10).

Many viruses, including SARS-CoV-2, have developed strategies to antagonize the autophagy pathway and thus escape host cell immunity^38,39^. Small GTPase proteins ARF1 and ARF6 act in the early steps of the autophagosome formation, and recent studies postulated that ARF6 might be bound and inhibited by the SARS-CoV-2 protein NSP15^40,41^. We have found SMAP2, a GTPase activating protein interacting with the ARF1 and ARF6 proteins, to be upregulated in SARS-CoV-2^+^ spots (Figure 2f, Supplementary Table 10). By catalyzing the GTP hydrolysis of ARF1 and ARF6 and thus rendering them in an inactive state, an increase in SMAP2 expression would mean a plausible alternative mechanism for the SARS-CoV-2 virus to block the autophagy pathway and promote viral replication and dissemination.

Previous work highlighted the NFκB pathway as a central signaling pathway in COVID-19 pathogenesis and the initiator of the so-called cytokine storm, characteristic of the disease^42,43^. In line with studies describing the direct induction of the NFκB pathway components by the SARS-COV-2 virus^44,45^, we found NFKB2 and NFKBIA, along with CXCL9, CCL17 and CCL21, to be upregulated in SARS-CoV-2^+^ spots (Figure 2f-g, Supplementary Table 10), and a study showed that the ORF7 protein of the SARS-CoV-2 virus induced these genes in an NFκB-dependent manner^46^. Previous studies proposed CCL17 as a potential predictive biomarker to distinguish between mild/moderate and severe/critical disease^47^, and CXCL9 to be part of a biomarker panel associated with mortality in patients with COVID-19^48^. Notably, we identified certain complement factors (C1QB, CFD, C7) and interferon response genes (IFI6, ISG15) to be upregulated in COVID-19 lungs compared to control lungs (Supplementary Table 6), in line with previous studies^3,7,49^; however, these genes were downregulated in the SARS-CoV-2^+^ spots of the infected COVID-19 lungs (Figure 2f, Supplementary Table 10), which points to spatially localized differences in host response to the virus as previously observed in terms of the interferon response genes^4^. A study proposed that the SARS-CoV-2 virus can direct a reduction in IFN response^50^, such as through NSP3 cleavage of ISG15^51^, which in turn might affect the subsequent activation of the complement system^52,53^.

In addition, we detected a downregulation of certain immunoglobulin genes (IGKC, IGKV4-1, IGHA1, IGHG2) and extracellular matrix components (FBLN, COL1A2, COL3A1, BGN, COL1A1, SPP1) in the SARS-CoV-2^+^ spots in the diseased COVID-19 tissue samples, which may be explained by the viral infection preceding both the extensive plasma cell infiltration and fibroblast activation in time (Figure 2f, Supplementary Table 10). It is also possible, however, that areas closely resembling the healthy state of the lungs are more permissive for the replication of viral components. Finally, we found several AT2 (SFTPB, SFTPC, MUC1, SLC34A2), AT1 (GPRC5A, AGER) and the alveolar endothelial cell marker AQP1 to be downregulated in the SARS-CoV-2^+^ spots (Figure 2f, Supplementary Table 10). These differences can either represent functional impairment or increased apoptosis of alveolar epithelial cells, which are known to be the primary cellular targets of the SARS-CoV-2 in the lungs^3,49^.

## Discussion

Future work on SARS-CoV-2-human spatially resolved interactions could utilize the method proposed here to explore how the different SARS-CoV-2 genes regulate host gene expression through the colocalization of specific SARS-CoV-2 genes with the human transcriptome. Furthermore, additional probes that indicate if SARS-CoV-2 is actively replicating by targeting the negative strand of the viral genomic RNA (gRNA) could be developed. In terms of the general outlook on spatially resolved host-pathogen interactions, limitations of our proposed approach include the requirement of previous knowledge of the pathogen transcriptome of interest to develop targeted probes, the inability to detect different human RNA splice variants, the lack of capturing human non-coding RNA groups that may have important regulatory functions, and the inability to detect new viral variants since the viral RNA is not directly sequenced. However, probes targeting specific host RNAs of interest could be developed to overcome some of these shortcomings.

In conclusion, the proposed method enables insights into highly localized host response to pathogen infection within the spatial context of the tissue microenvironment at the whole-transcriptome level in an unbiased and high-throughput manner. The method has the potential to be applied to other human pathogens with the development of targeted probes and thus examine the interplay between host and pathogen across the multitude of human infectious diseases. Overall, our approach opens the door to new possibilities of studying infectious disease at a large scale by exploring multiple transcriptomes in a single experiment.

## Author contributions

S.G. and O.B. conceived and designed the project. T.V., T.B., A.K. performed sample collection. H.S. performed spatial transcriptomics and *in situ* sequencing experiments. E.L. performed RNAscope experiments. A.J. provided assistance with spatial transcriptomics experiments. A.N. provided assistance with *in situ* sequencing experiments. A.H. designed SARS-CoV-2 probes. E.B. performed read alignments. Y.M. performed computational analysis of the spatial transcriptomics data and colocalization analysis with supervision of S.G. J.W. designed the computational approach for spatial transcriptomics, RNAscope, and *in situ* sequencing validation and performed the analysis with supervision of S.G. E.L. performed image annotation. H.S., E.L., H.D., O.B. and S.G. interpreted the results. H.S. prepared figures with input from S.G. S.G. and O.B. supervised and guided the project. H.S., E.L., O.B. and S.G. wrote the manuscript with input from Y.M. and J.W. All the authors read and approved the manuscript.

## Acknowledgements

This work was supported by the Swedish Research Council as Formas grant 2017-01066 and Vetenskapsrådet grant 2020-04864 to S.G. O.B. was supported by the Center for Regenerative Therapies Dresden, the Karolinska Institute, the Swedish Research Council, the Ragnar Söderberg Foundation, the Åke Wiberg Foundation, and the LeDucq Foundation. We thank 10x Genomics for support with instruments and/or lab reagents and technical advice (Patrick Roelli and Iván Hernández). We thank Karl Frontzek for help with providing lung samples.

## Methods

### Patient selection, sample collection and processing

Collection of postmortem samples from lung tissue was performed at the 2^nd^ Department of Pathology, Semmelweis University (Budapest, Hungary) and the University Hospital Zurich (Switzerland). Autopsy cases were selected from patients who were hospitalized because of COVID-19 infection and died at the local clinical departments of the universities. Criteria for selection were: premortem positive (COVID-19 cases) or negative (control cases) SARS-CoV-2 PCR test, lack of malignancy of the lung, closed clinical documents and less than 24 hours as a postmortem interval (PMI). Autopsy was done in harmony with the World Health Organization’s (WHO) recommendation for autopsy of COVID-19 cases^54^. The biopsies were fixated in formaldehyde (4%) and then went through a dehydration process overnight. Dehydrated samples were embedded into paraffin blocks and were stored at 4°C until sectioning. The use of tissue specimens collected at Semmelweis University in this study was approved by the Hungarian Scientific Research Ethics Committee (ETT TUKEB IV/3961-2/2020/EKU). Samples and data were managed anonymously. At the University Hospital Zurich, small quantities of bodily substances removed in the course of an autopsy were anonymized for research purposes without consent, in the absence of a documented refusal of the deceased persons. In accordance with the Swiss Federal Act on Research involving Human Beings, this study did not require institutional board approval. Subsequent experiments were approved by the Swedish Ethical Review Authority (2010/313-31/3, 2018/689-32). Relevant clinical parameters of the patients included in this study are summarized in Supplementary Table 2.

### Sample selection - Evaluating RNA Quality

Total RNA was extracted from each formalin-fixed paraffin-embedded (FFPE) sample block with the RNeasy FFPE kit (Qiagen, Cat. No. / ID: 73504) following the manufacturer’s instructions (deparaffinization was performed using xylene (#28975.291 VWR) and 96% EtOH (#20823.290 VWR) or 100% EtOH (#1.00983.1000 VWR)). The concentration of extracted total RNA was determined with the RNA HS Qubit assay (Thermo Fisher Scientific) following the manufacturer’s instructions. Total RNA was diluted to between 2-5ng and RNA fragment length assessed using the Agilent RNA 6000 Pico Kit following the manufacturer’s instructions. The RNA quality of the sample was evaluated by the DV200 measurement (percentage of RNA fragments longer than 200 nucleotides) as specified in the Visium Spatial Gene Expression for FFPE – Tissue Preparation Guide^55^. Samples with a DV200 greater than 40% were selected for Visium FFPE, RNAScope and *in situ* sequencing.

### SARS-CoV-2 probe design

SARS-CoV-2 probes were designed as described^56^, with probes designed based on the reference transcriptome Sars_cov_2.ASM985889v3, Ensembl build 101 (https://covid-19.ensembl.org/Sars_cov_2/Info/Index). Probes were designed to target SARS-CoV-2 genes: Surface glycoprotein (S), Envelope protein (E), Membrane glycoprotein (M), ORF1ab, ORF3a, ORF7a, ORF7b, ORF8, Nucleocapsid phosphoprotein (N), and ORF10 (10x Genomics) (Supplementary Tables 1,11-12).

### Spatial Transcriptomics

Consecutive 5μm tissue sections from each sample were placed onto Visium Spatial Gene Expression slides (PN: 2000233, 10X Genomics) and stored overnight in a desiccator^55^. 5μm consecutive sections to the sections used for Visium FFPE were placed onto Superfrost Plus microscope slides (#631-9483, VWR) and stored at 4°C until used for RNAScope and *in situ* sequencing. Deparaffinization, Hematoxylin and Eosin staining, and decrosslinking were performed as specified in the the Visium Spatial Gene Expression for FFPE – Deparaffinization, H&E Staining, Imaging & Decrosslinking Demonstrated Protocol^57^. Spatial gene expression profiling of RNA from FFPE lung samples was performed by following all steps in the Visium Spatial Gene Expression Reagent Kits for FFPE User Guide^20^ with the modifications: For COVID-19 samples (see Supplementary Table 2), four 5μm consecutive sections per patient sample tissue FFPE block were placed on Visium Spatial Gene Expression slides (PN: 2000233, 10x Genomics). For step 1.1.g, Human whole transcriptome (WT) probes (10x Genomics) were added to two consecutive sections (technical replicates) with the Probe Hybridization Mix: 19.8μL Nuclease-free water, 77.0μL FFPE Hyb Buffer, 6.6μL LHS Human WT probes, and 6.6μL RHS Human WT probes, per sample. Human WT and spike-in custom probes targeting SARS-CoV-2 genes (10X Genomics) were added to the remaining two consecutive sections (technical replicates) with the Probe Hybridization Mix: 14.5μL Nuclease Free water, 77.0μL FFPE Hyb Buffer, 6.6μL LHS Human WT probes, and 6.6μL RHS Human WT probes, 2.6μL LHS viral probes, and 2.6μL RHS viral probes, per sample. For control patient samples, two consecutive sections (technical replicates) were processed as described for the COVID-19 samples, with adding Human WT and SARS-CoV-2 spike-in probes to all sections. For step 4.1.d, qPCR (Bio-Rad) step 4 was run for a total of 30 cycles. For step 4.2.d, the Sample Index PCR was performed with 15 cycles for 1C, 15-16 cycles for 3C, 18-19 cycles for 2C, 16 cycles for 4nC, and 18 cycles for 5nC. After step 4.4, the concentration of sequence libraries were determined with 2μL of each sample run with the dsDNA HS Qubit assay (Thermo Fisher Scientific).

### Spatial Transcriptomics Hematoxylin & Eosin Imaging

Hematoxylin & Eosin brightfield images were acquired with a Zeiss Axiolmager.Z2 VSlide Microscope using the Metasystems VSlide scanning system with Metafer 5 v3.14.179 and VSlide software. The microscope has an upright architecture, uses a widefield system, and a 20X air objective with the numerical aperture (NA) 0.80 was used. The camera was a CoolCube 4m with a Scientific CMOS (complementary metal-oxide-semiconductor) architecture and monochrome with a 3.45 x 3.45 μm pixel size. All brightfield images were taken with a Camera Gain of 1.0 and an Integration Time/Exposure time of 0.00011 seconds.

### Spatial Transcriptomics Sequencing

Sequencing libraries were pooled and diluted with Elution Buffer (EB) to a final concentration of 10nM, using a target sequencing depth of 50,000 mean read pairs/spot to determine the dilution for each sample^20^. After sample pooling, pooled library concentrations were checked with qPCR (Bio-Rad) before loading into the sequencer. Libraries were sequenced on an Illumina NovaSeq 6000 with paired-end, dual indexed sequencing run type and parameters following those specified in the Visium Spatial Gene Expression Reagent Kits for FFPE User Guide sequencing instructions^20^ [R1: 28 cycles, R2S: 50-52 cycles], with a spike-in of PhiX at 1% concentration, except one sample, 3C, was run with R2S: 75 cycles.

### *In situ* sequencing (ISS)

Optimal RNA integrity and assay conditions were assessed using *MALAT1* and *RPLP0* housekeeping genes only using the HS Library Preparation kit for CARTANA technology (part of 10x Genomics) and following manufacturer’s instructions on 5μm tissue sections from representative sample 1C. Since the control probes test showed positive and expected results, *in situ* sequencing was then performed on two 5μm consecutive sections from sample 1C and one consecutive section from each control sample (4nC and 5nC). Superfrost Plus microscope slides (#631-9483, VWR) containing 5μm tissue sections were stored at 4°C until processing. FFPE sections were baked for 1 hour at 60°C to partially melt paraffin and increase tissue adherence. Next sections were deparaffinized using xylene for 2×7 minutes followed by an EtOH gradient to remove xylene and rehydrate the sections. Sections were then permeabilized using citrate buffer pH 6.0 (C9999 Sigma Aldrich) for 45 minutes at 95°C. For library preparation, chimeric padlock probes (targeting directly RNA and containing an anchor sequence as well as a gene-specific barcode) for a custom panel of SARS-CoV-2 S and E genes were hybridized overnight at 37°C, then ligated before the rolling circle amplification was performed overnight at 30°C using the HS Library Preparation kit from CARTANA technology and following manufacturer’s instructions. All incubations were performed in SecureSeal™ chambers (Grace Biolabs). For tissue section mounting, Slow Fade Antifade Mountant (Thermo Fisher) was used for optimal handling and imaging. Quality control of the library preparation was performed by applying anchor probes to simultaneously detect all rolling circle amplification products from all genes in all panels. Anchor probes are labeled probes with Cy5 fluorophore (excitation at 650 nm and emission at 670 nm). All samples passed the quality control l and were sent to CARTANA (part of 10x Genomics), Sweden, for a single cycle *in situ* barcode sequencing, imaging and data processing. Briefly, adapter probes and a sequencing pool (containing 4 different fluorescent labels: Alexa Fluor® 488, Cy3, Cy5 and Alexa Fluor® 750) were hybridized to the *in situ* libraries to detect SARS-CoV-2 gene-specific barcodes. This was followed by multicolor epifluorence microscopy, scanning the whole area and thickness of the tissues. Raw data consisting of 20x magnification images from 5 fluorescent channels (DAPI, Alexa Fluor® 488, Cy3, Cy5 and Alexa Fluor® 750) and individual z-stacks, were flattened to 2D using maximum intensity projection with a Nikon Ti2 Nikon Ti2 (software NIS elements) utilizing Zyla 4.2 camera. After image processing, which includes image stitching, background filtering and a sub-pixel object registration algorithm, true signals were scored based on signal intensities from individual multicolor images. The results were summarized in a csv file and gene plots were generated using MATLAB.

### RNAscope assay and imaging

RNAscope assay was performed on lung 5 μm FFPE sections on Superfrost Plus microscope slides (#631-9483, VWR) cut from depths consecutive to the sections mounted on Visium slides. The slides were baked in a dry oven for 1 h at 60 °C and then deparaffinized in xylene (2x 5 min) and absolute ethanol (2x 1 min) at room temperature. After drying, the sections were incubated in RNAscope Hydrogen Peroxide for 10 minutes at room temperature, followed by washing steps (2x) in distilled water. Target retrieval was performed using a 1x RNAscope Target Retrieval Reagent for 15 minutes, at a temperature constantly kept above 99 °C in a hot steamer. The slides were then rinsed in distilled water, incubated in absolute ethanol for 3 minutes and dried at 60 °C. After creating a hydrophobic barrier, the slides were left to dry overnight. The second day, the sections were incubated in RNAscope Protease Plus solution for 30 min at 40 °C, followed by washing in distilled water. RNAscope V-nCov2019-S probe, RNAscope Positive Control probe (Hs-PPIB) and RNAscope Negative Control Probe (DapB) were hybridized to separate sections for 2h at 40 °C, then the slides were washed twice for 2 minutes in 1x Wash Buffer. The probe-specific signal was developed with an RNAscope 2.5 HD Detection Reagent – RED kit. Sequential hybridization of amplification reagents AMP1-4 happened at 40 °C for 30-15-30-15 minutes, while AMP5 and AMP6 were applied at room temperature for 1 hour and 15 minutes, respectively, with two washing steps in 1x Wash Buffer after each incubation period. For signal detection, each section was incubated for 10 min at room temperature in 120 ul RED Working Solution, consisting of Fast RED-B and Fast RED-A reagents in a 1:60 ratio. All the protease digestion, probe hybridization, signal amplification and signal detection steps were performed in a HybEZ Humidity Control Tray, which was either placed into a HybEZ Oven for the 40 °C incubation steps or kept at room temperature. Following two washing steps in tap water, the slides were counterstained with 50% Gill’s Hematoxylin staining solution for 2 min at room temperature, thoroughly rinsed with tap water, then soaked in 0.02% Ammonia water bluing solution and finally washed again in tap water. The slides were then dried completely at 60 °C and then quickly dipped into xylene before mounting them with VectaMount Permanent Mounting Medium. The RNAscope signal was imaged and evaluated with a Leica DM5500B microscope with a HC PL APO 20x/0.70 DRY objective, using Extended Depth of Field (EDoF) imaging in the Leica Application Suite X (LAS X) software platform.

### Spatial Transcriptomics - Data Processing

#### Count matrices generation

The gene expression matrices were generated by spaceranger (version 1.3.0) ‘count’ (standard settings set except -- *no-bam*). The transcriptome reference was custom made from spaceranger ‘mkref’ using Human reference dataset (GRCh38 Reference - 2020-A), and SARS-CoV-2 genome assembly (ASM985889v3). The Human Probe Set from 10x Genomics (Visium Human Transcriptome Probe Set v1.0) with 10x Genomics custom probes for SARS-CoV-2 probes appended to it, was used as the probe set reference in spaceranger ‘count’.

#### Quality Control

The filtered count matrices (filtered_feature_bc_matrix.h5), and tissue images from spaceranger output were analyzed in R using the Load10X_Spatial function available in Seurat (version 4.0.4)^58^. The filtered count matrices were separated into human count data, and SARS-CoV-2 count data matrices. Spot level filtering was performed on the human count matrices to keep spots with at least 400 genes, 500 UMIs, and a novelty score of 0.87. Gene level filtering was applied to omit genes that did not appear in at least 1 spot. These count matrices were also filtered for Hemoglobin gene counts (Supplementary Table 1). SARS-CoV-2 count matrices were normalized by dividing the SARS-CoV-2 gene UMI counts by the number of probes used to target the respective gene. 1 SARS-CoV-2 UMI was detected from two different sections, one control and one COVID-19, that did not have SARS-CoV-2 probes added and was considered as background signal.

#### Clustering Analysis

The Seurat SCTransform function was applied to normalize the individual filtered count matrices, and integrated in Seurat using SelectIntegrationFeatures, and IntegrateData. Principal Component Analysis (PCA), and UMAP was applied using 50 principal components, and 35 were further used in downstream analysis, and clustering. Batch effects were addressed, and removed using RunHarmony (group.by.vars as slide ID, and 25 iterations) applied on the PCA-computed matrix ^59^. Clustering was applied at a resolution of 0.4.

#### Differential Gene Expression

Differentially expressed (DE) genes were found using ‘FindMarkers’ in Seurat, with default settings on the SCT normalized matrix, except min.cells.group set to 2 to include at least 2 spots from each group. Both upregulated and downregulated DE genes were identified, with an adjusted p-value of 0.005. Cell-type specific annotation of the DE genes was performed manually, by using the Human Single Cell Atlas^60^, PanglaoDB^61^, and recently published single cell transcriptomic data of the human lung^3,49^ as main resources.

#### Colocalization analysis

For the colocalization analysis, a direct spot-level comparison within the COVID-19 sections was performed. The DE genes distinguishing SARS-CoV-2^+^ spots from SARS-CoV-2^-^ spots were obtained as described in the Methods section “Differential Gene Expression” with an additional filter of average logFC +/- 1.0.

#### Validation by RNAScope

RNAScope and ST images were manually aligned with Adobe Photoshop 2022. The RNAScope chromogenic detection of the S gene with FastRed was used to distinguish RNAScope signal from lung pigmentation and tar deposits. All dots of chromogenic red signal were considered as positive SARS-CoV-2 S gene signal, since the majority of signal was above 1 dot per 10 nuclei area, in line with how others assessed RNAScope signal in SARS-CoV-2 viral low samples^3,4,62^. RNAScope was considered as the gold standard for comparison to the ST signal. The number of ST spots where the SARS-CoV-2 S gene was detected, and where the RNAScope S gene signal was also obtained, was calculated. To adjust for the use of consecutive sections for ST and RNAScope experiments, the agreement of ST and RNAScope in 200×200 *μm^2^* block areas were evaluated. Since a manual annotation of sample 1C was in close agreement with the computational approach, the computational approach to calculate the specificity of the SARS-CoV-2 S gene detection by ST was used.

The computational validation was performed as follows: the RNAScope signal was detected with an ad hoc Matlab (version R2021b) algorithm, which is specified in the next section “Automatic detection of RNAScope signal”; then both the binary ST and RNAScope signal images were aligned and binned into 200×200 *μm^2^* blocks (Supplementary Figures 3-4). Each block in an RNAScope/ST signal image was regarded as an observation (those blocks that contain no tissue area were regarded as no observation and were excluded from any further analysis and counting). The specificity of our method to capture the SARS-CoV-2 expression was calculated by considering the RNAScope approach as the groundtruth and as follows:

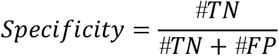

Where the number (#) of True Negatives (TN) was defined as the number of blocks containing neither RNAScope nor ST signals and the number of False Positives (FP) as the number of blocks containing only ST but no RNAScope signals.

#### Automatic detection of RNAScope signal

RNAScope signals were detected with a chromatical analytic method. First, the original RGB image was transformed into the Hue-Saturation-Value (HSV) format, where the bright regions in the hue channel correspond to the RNAScope signals in the original histological image. The brightest regions became the foreground by thresholding the hue value of the image. Morphological post processing steps were performed to refine the shape of the signal regions, the details of which are available in the code (see “Code availability”). The pixels whose hue was over 0.85, saturation over 0.25, and value over 0.40 were recognized as signal candidates. After performing a morphological opening operation, the collection of signal candidates were output as final RNAScope signals. Supplementary Figure 5 displays the original tissue subimage, the hue channel, and the RNAScope signal subimage after the thresholding.

#### Validation by ISS

ISS consecutive section images and ST images for sample 1C were manually aligned with Adobe Photoshop 2022. Due to the use of non-consecutive sections, there was ~300 μm in between the ST and ISS sections. The agreement between E and S gene signals for ST and ISS in block areas of 200μm was evaluated using the same computational approach as used for the RNAScope validation.

## Data availability

Raw sample sequences will be available with controlled access on the European Genome-Phenome Archive (EGA). Processed gene count matrices, related metadata, corresponding ST tissue microscopy images, and images used in the Validation analysis are available on Mendeley dataset under the Reserved DOI: 10.17632/xb2w7xvs2b.1. ISS images for SARS-CoV-2 S and E genes can also be downloaded from Mendeley dataset with Reserved DOIs 10.17632/gwjk2cxsf4.1. RNAscope images for SARS-CoV-2 S gene are available on FigShare project ID 134597 which is currently private. RNAScope images for positive and negative control probes are available upon request.

## Code availability

Scripts to generate the count matrices and all related R scripts used in the clustering, differential expression, colocalization analysis, and the program for the computational validation can be accessed from our github repository, DualST]Study.

