## Supplementary Information for "Dual spatially resolved transcriptomics for SARS-CoV-2 host-pathogen colocalization studies in humans"

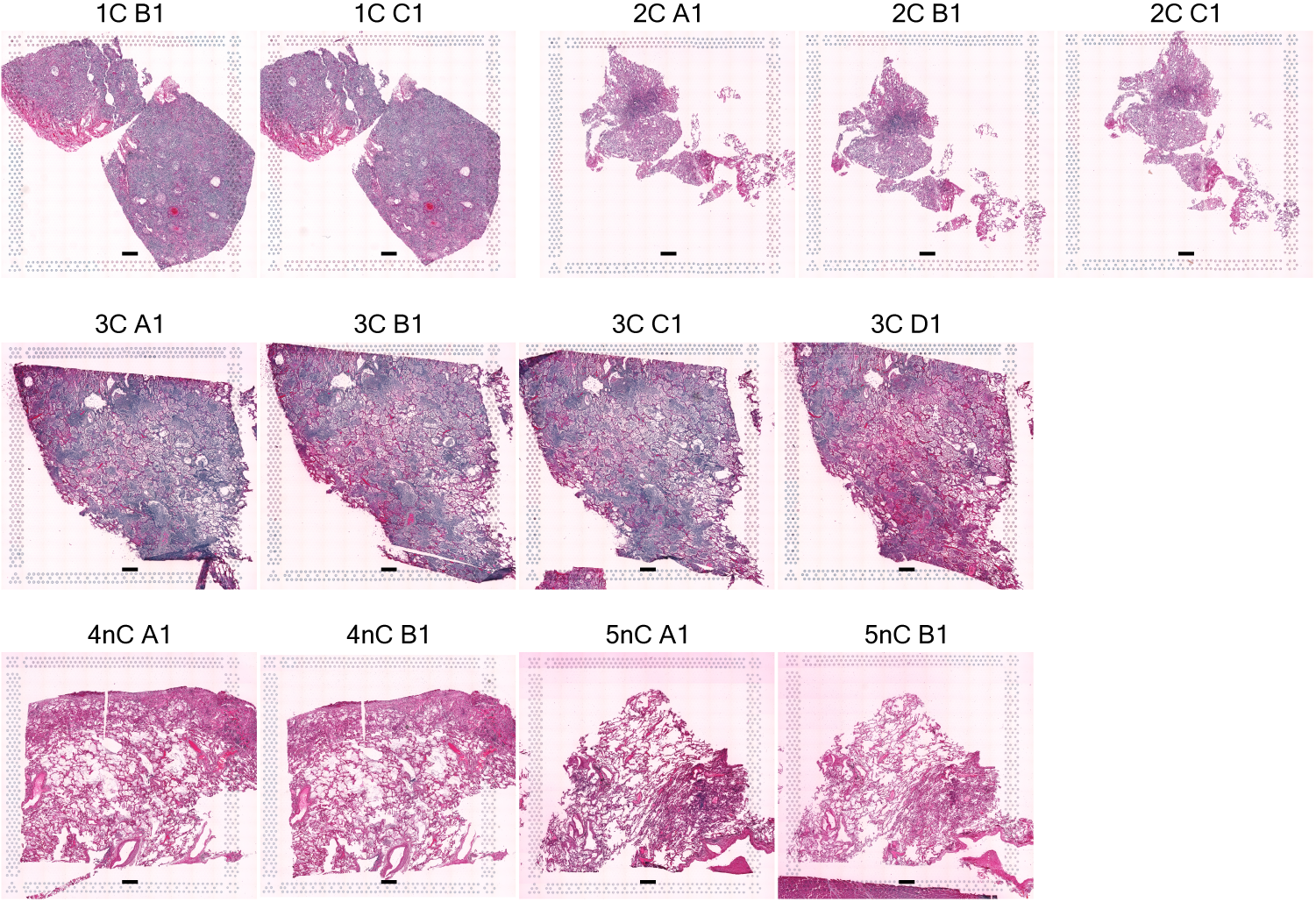

**Supplementary Figure 1. Study sample sections.** The 13 tissue sections across 5 patient lung samples, 3 COVID-19 samples (1C, 2C, 3C) and 2 control samples (4nC, 5nC), used for ST in this study. Scale bars are 500µm.

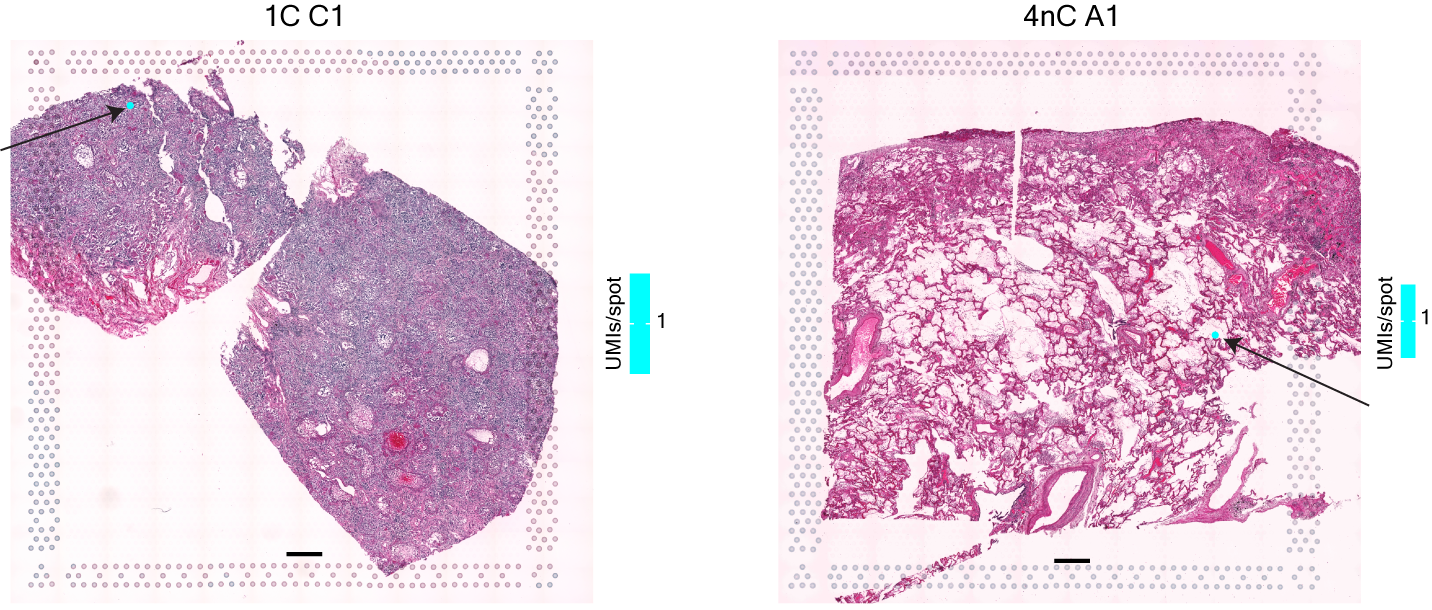

**Supplementary Figure 2. Background SARS-CoV-2 detection.** 1 UMI count across all assayed SARS-CoV-2 genes was detected in two tissue sections, a COVID-19 section with no SARS-Cov-2 probes added (1C C1) and a control section (4nC A1). Scale bars are 500µm.

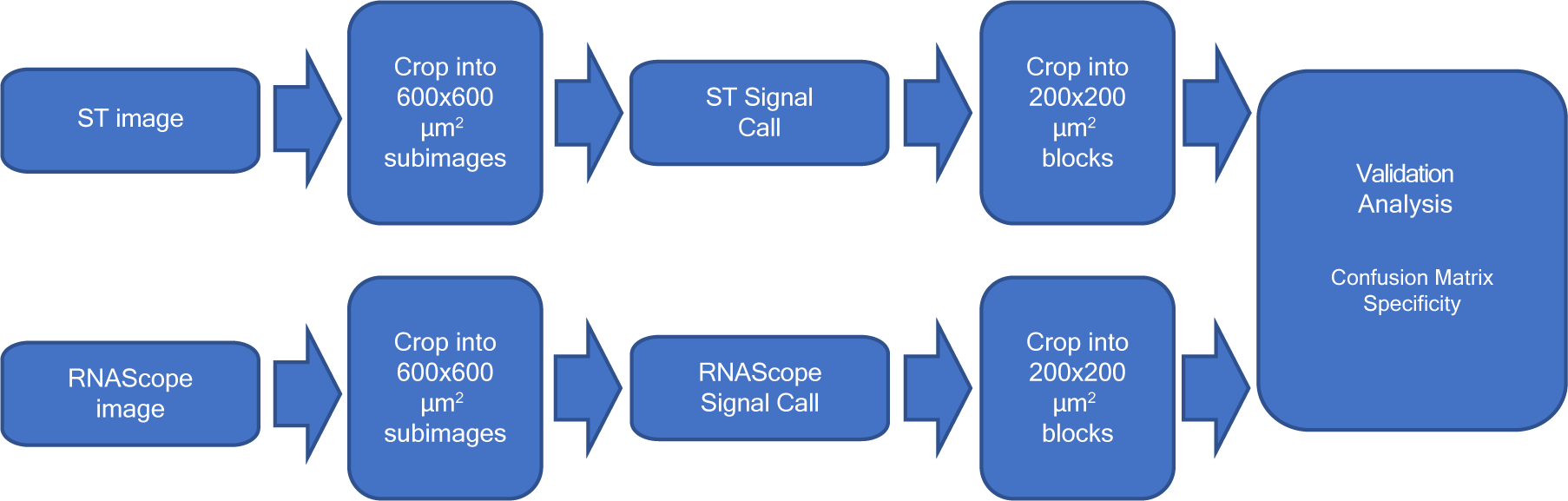

**Supplementary Figure 3.** **RNAScope Validation Workflow.** The RNAScope validation workflow consists of two independent and similar procedures, which process the RNAScope or ST image signals respectively, and compare the two procedure outputs. In each RNAScope/ST procedure, the original histological image is cropped into median sized subimages. The RNAScope/ST signals are detected by the corresponding method. Afterwards, the signal images are cropped into small blocks, which are utilized for the validation analysis.

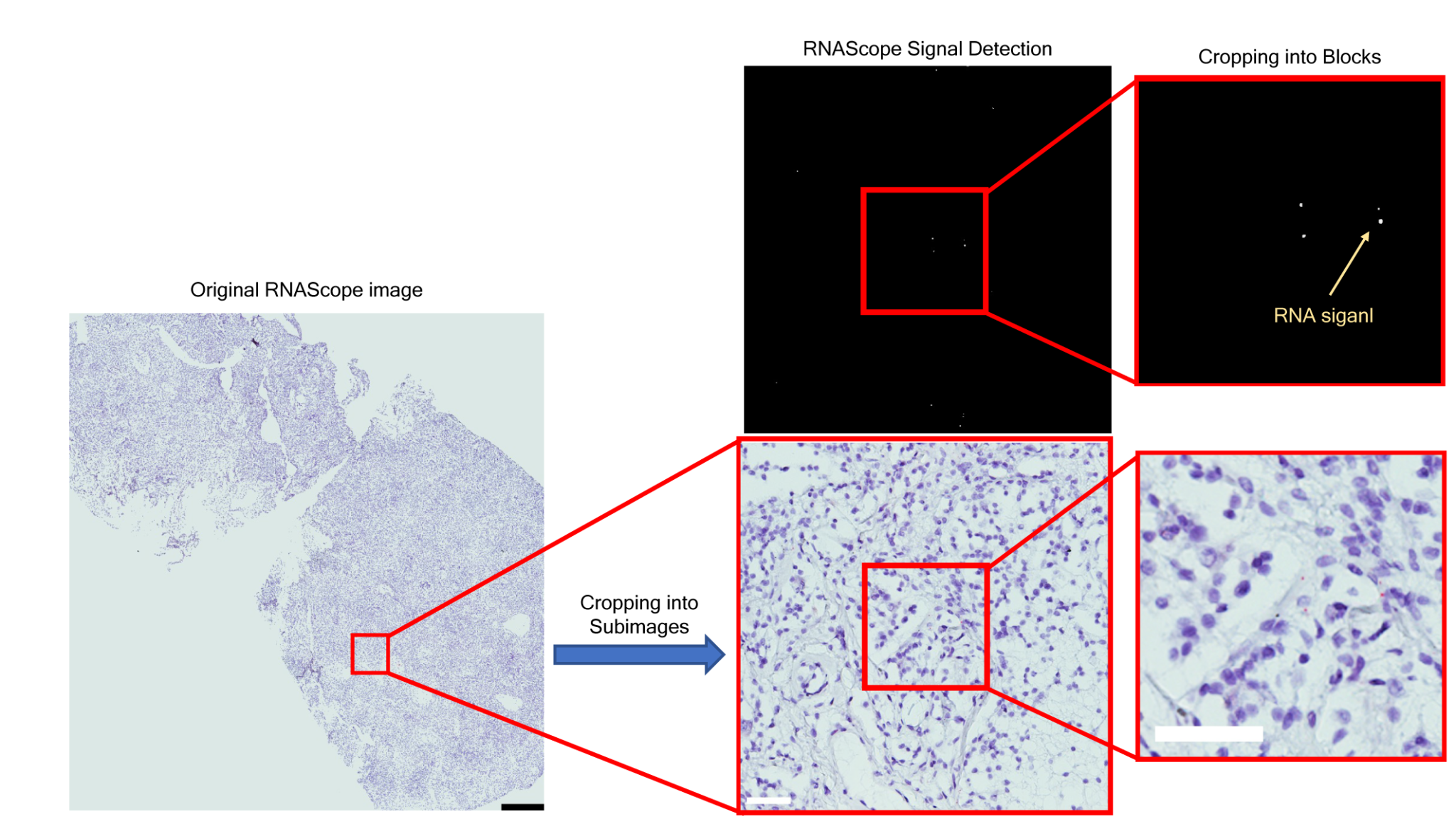

**Supplementary Figure 4. Workflow of RNAScope light-field image processing.** A visualization of the RNAScope signal processing procedure, where the RNAScope signal images are the same size as the corresponding light field images. The ST signal processing shares essentially the same procedure, except for the RNAScope/ST signal call. The black scale bar represents the length of 500µm and the white bars represent the length of 50µm.

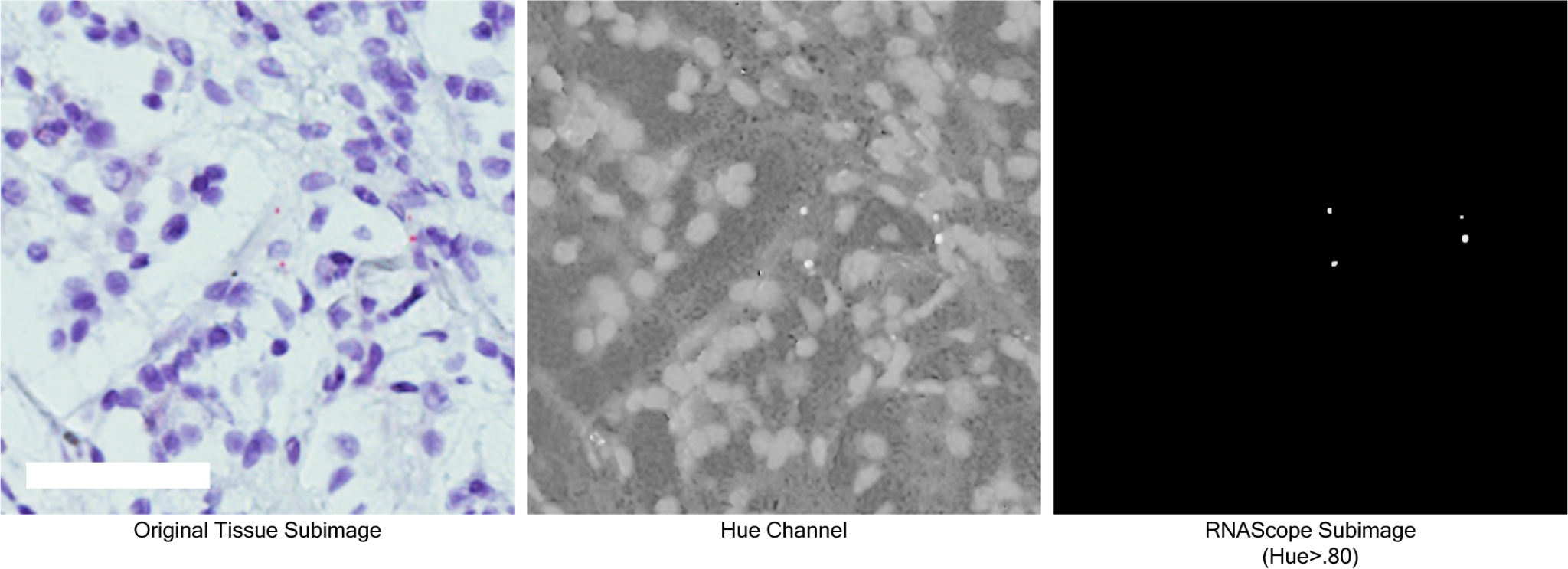

**Supplementary Figure 5**. **RNAScope Signal Call.** The original tissue subimage, the hue channel, and the RNAScope signal subimage after the thresholding, where the white scale bar represents the length of 50µm, and all the images are of the same size and resolution.

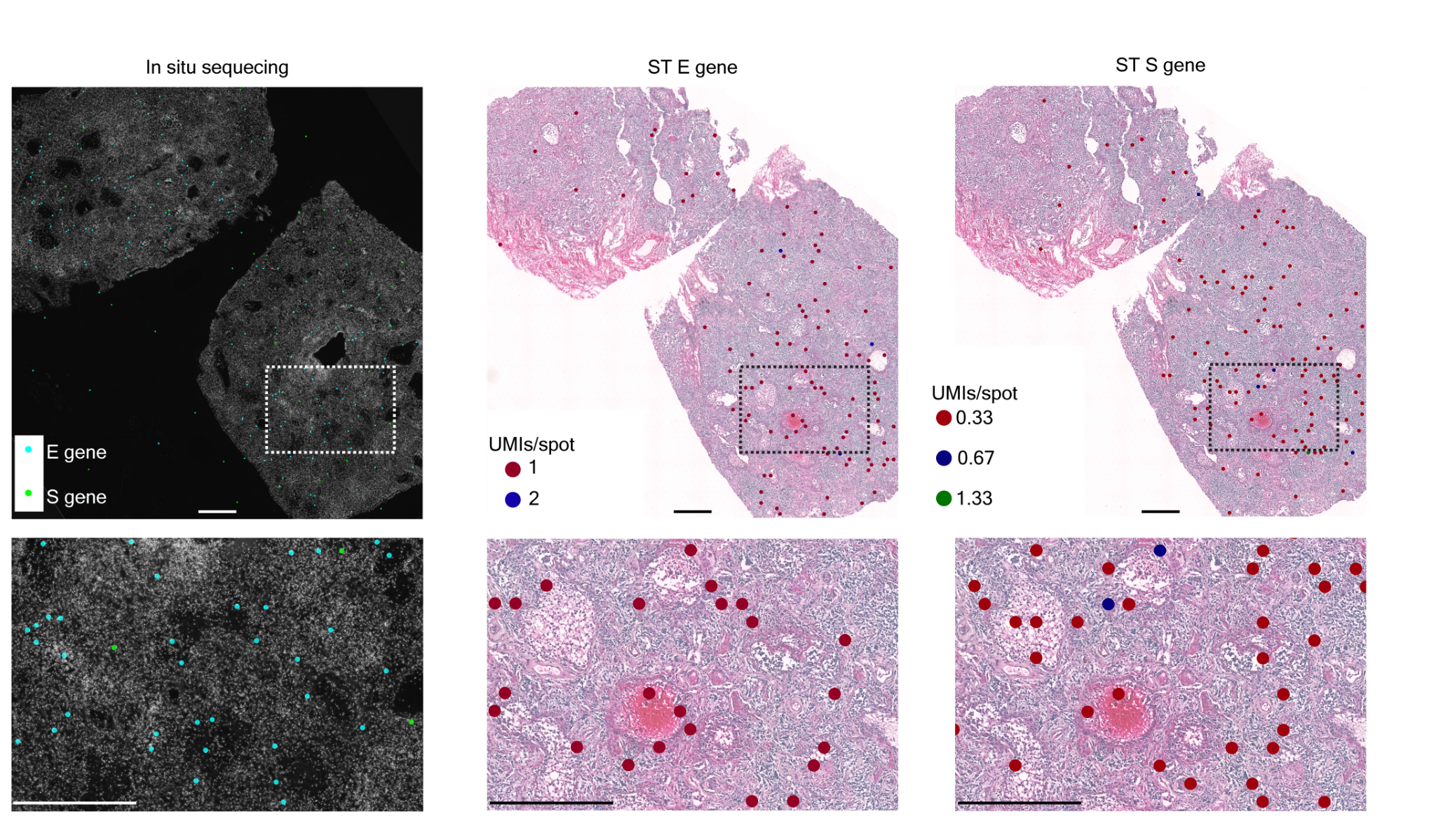

**Supplementary Figure 6. *In situ* sequencing validation of ST S & E gene signal.** Enlargement of Figure 1c showing the *in situ* sequencing (ISS) signal and ST signal for genes S and E (~300µm between ISS and ST sections). Scale bars are 550µm.

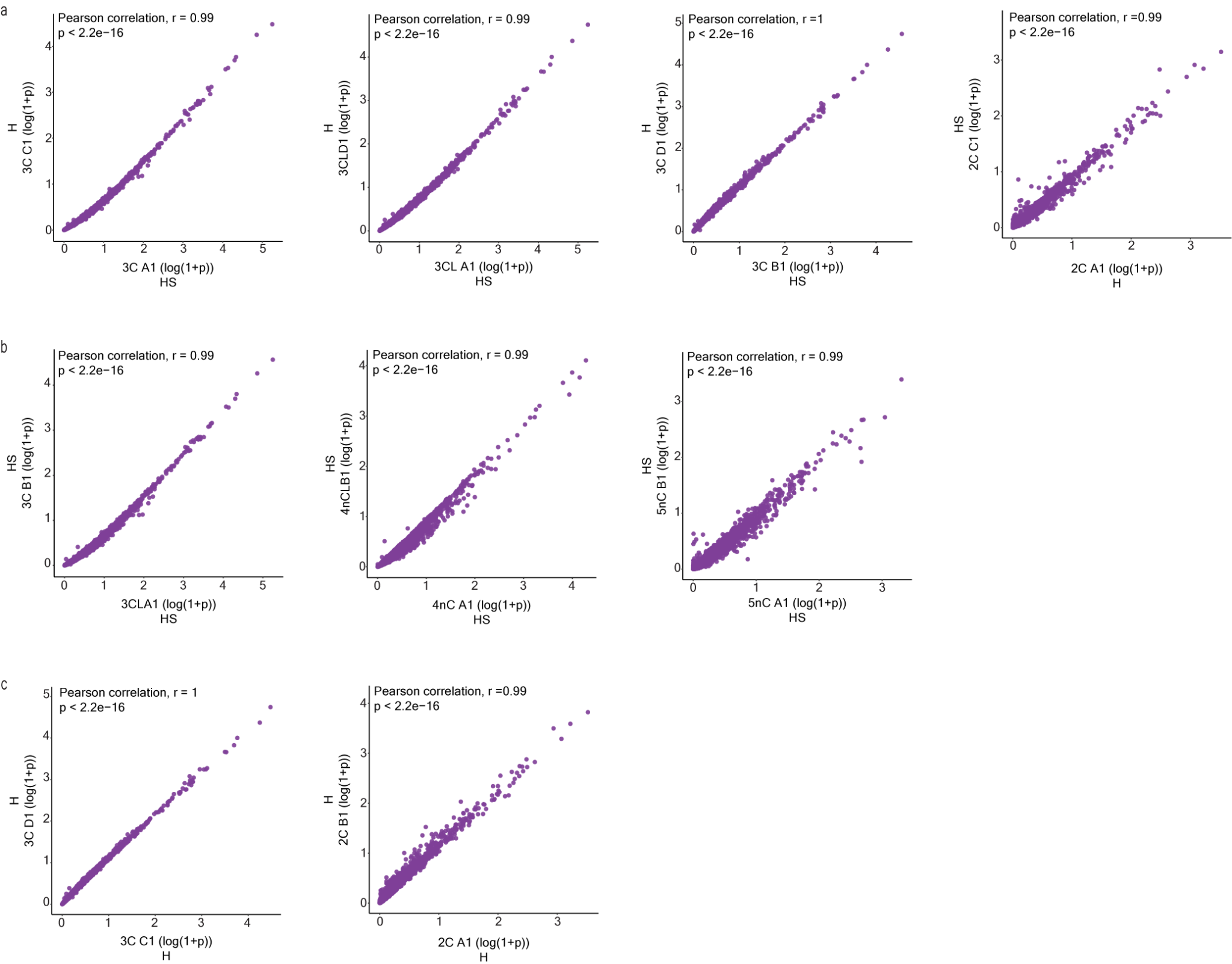

**Supplementary Figure 7**. **Highly reproducible capture of human transcriptome data.** Pearson correlation of average human gene expression between consecutive sections for each sample, with either (a) one section with human and SARS-CoV-2 probes added (HS) and the other with only human probes added (H), (b) both sections with human and SARS-CoV-2 probes added (HS), or both sections with only human probes added (H). P-value < 2.2e-16.

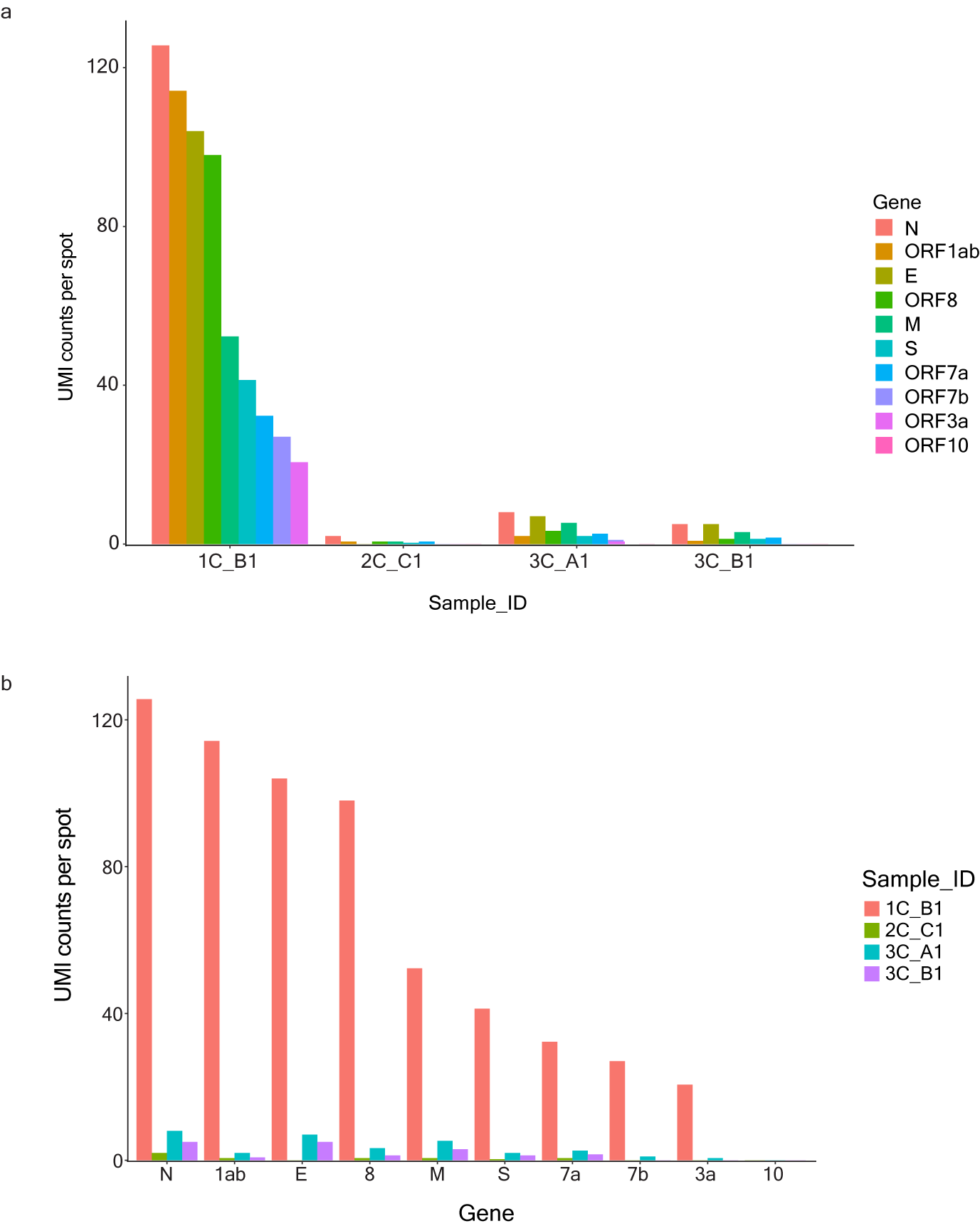

**Supplementary Figure 8. SARS-CoV-2 gene distributions for each sample section assayed.** (a) Total UMI counts per spot of each SARS-CoV-2 gene for each COVID-19 sample section. (b) Total UMI counts per spot of each SARS-CoV-2 gene across eachCOVID-19 sample section.

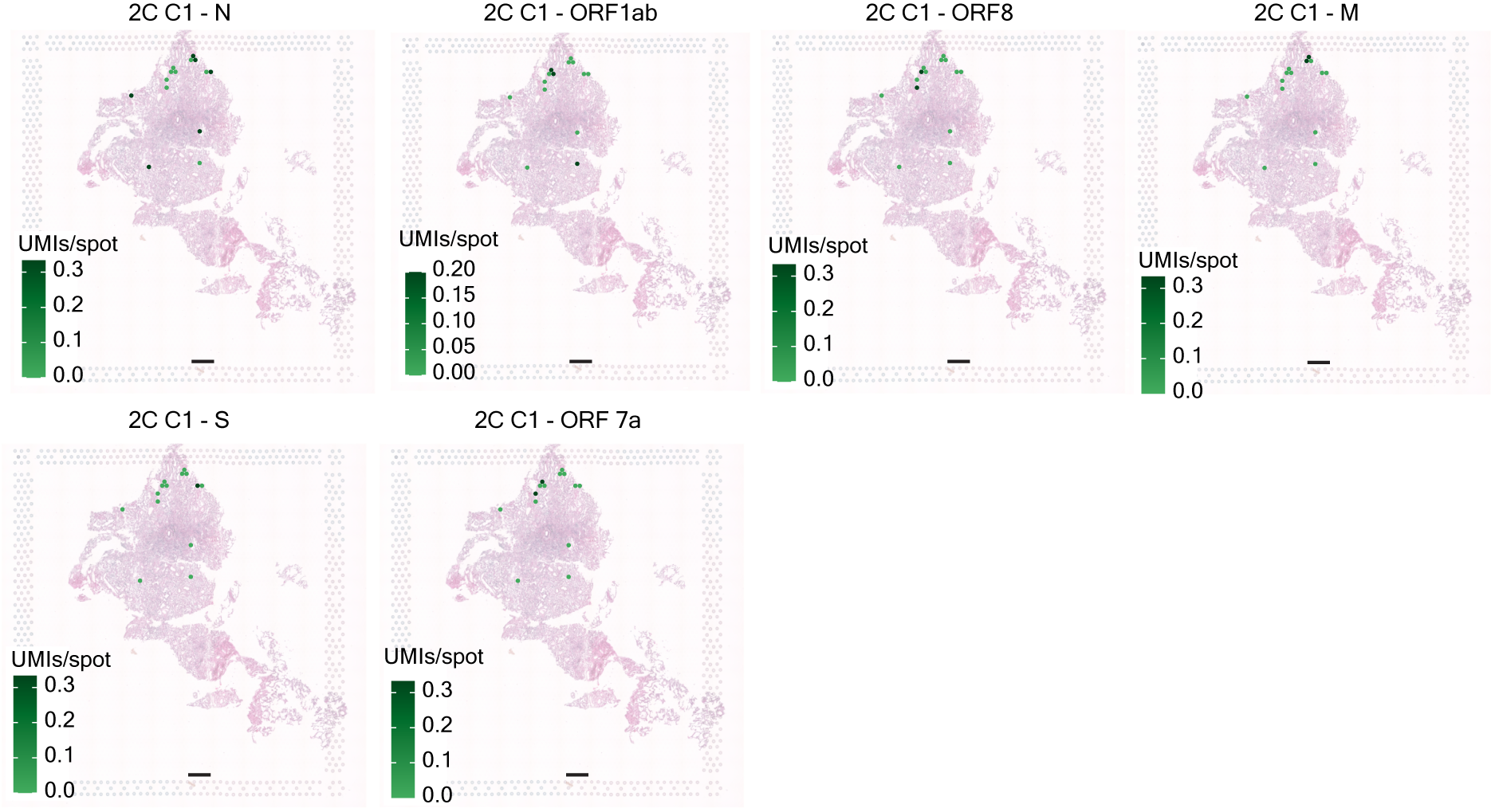

**Supplementary Figure 9. Spatial SARS-CoV-2 gene distributions across tissue sample section 2C C1.** The spatial distribution of UMI counts per capture spot of each SARS-CoV-2 gene across COVID-19 sample section 2C C1. Scale bars are 500µm.

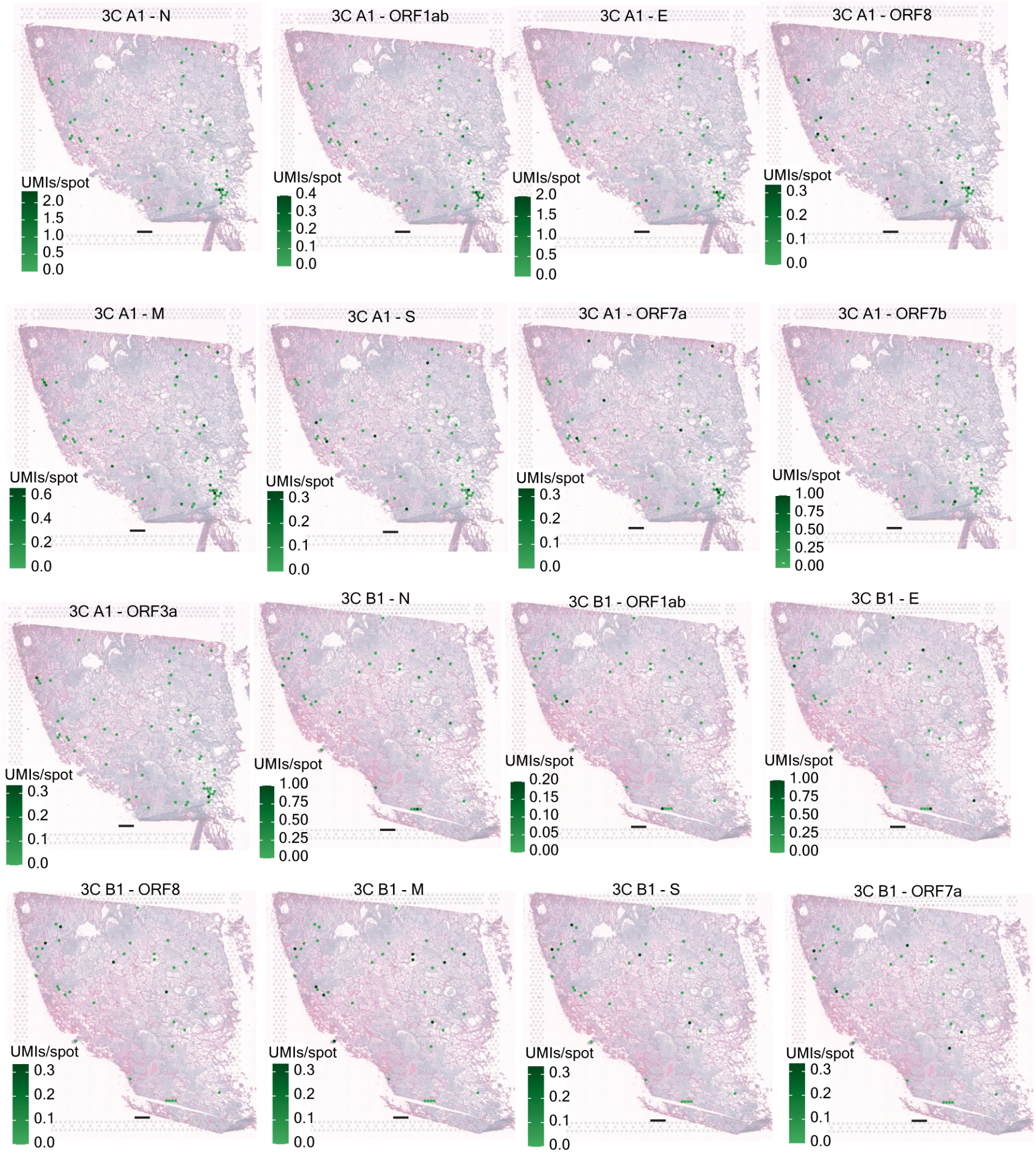

**Supplementary Figure 10. Spatial SARS-CoV-2 gene distributions across tissue sample 3C.** The spatial distribution of UMI counts per capture spot of each SARS-CoV-2 gene across COVID-19 sample sections 3C A1 and 3C B1. Scale bars are 500µm.

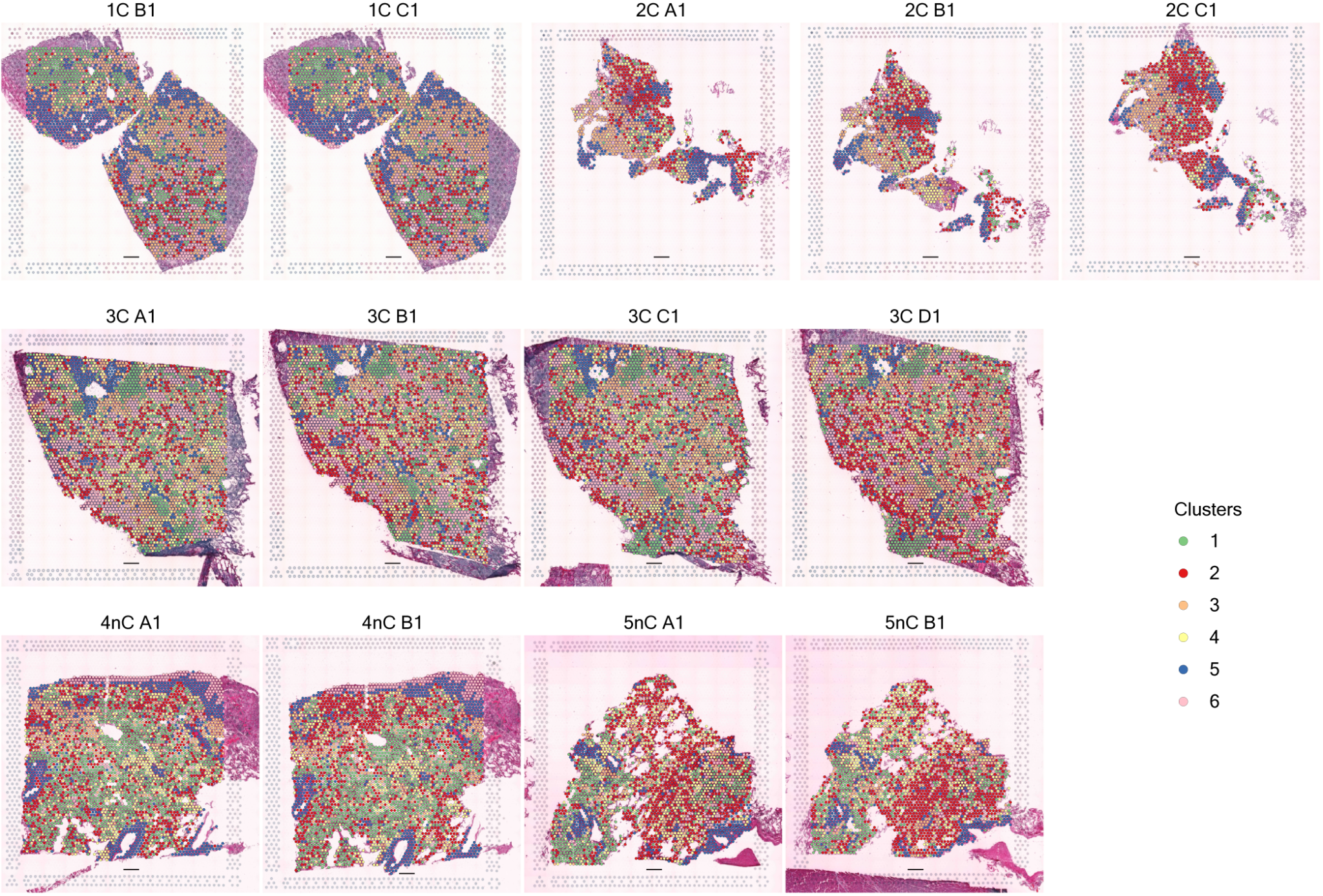

**Supplementary Figure 11. Clusters mapped onto the 13 ST tissue sections.** Spatial distribution of the clusters on COVID-19 and control sections. Scale bars are 500µm.

**Supplementary Table 1**. The human (16,688) and SARS-CoV-2 (10) gene transcripts targeted in the study.

| **Patient ID** | **Sample ID** | **COVID/Control** | **Survival after COVID diagnosis (days)** | **Postmortem interval (hours)** | **Fixation time (hours)** | **DV200** | **RIN** |
| --- | --- | --- | --- | --- | --- | --- | --- |
| P1 | 1C | COVID | 15 | 17 | 24 | 62 | 2.4 |
| P2 | 2C | COVID | 13 | 21 | 24 | 45 | 2.4 |
| P3 | 3C | COVID | 17 | 13 | 6 | 61 | 2.0 |
| P4 | 4nC | Control | - | 21 | o.n. | 63 | 2.2 |
| P5 | 5nC | Control | - | 15 | o.n. | 72 | 2.1 |

**Supplementary Table 2. Patient data.** Relevant clinical parameters of the 5 patients included in this study are summarized.

**Supplementary Table 3. Spatial transcriptomics sample section summary.** Sequence library information for each of the 13 patient tissue sections assayed with spatial transcriptomics (ST).

**Supplementary Table 4. Clustering differential expressed genes.** Differentially expressed (DE) genes for each spatial transcriptomic (ST) cluster.

**Supplementary Table 5. Cluster 5 subclustered differentially expressed genes.** Differentially expressed (DE) genes for each subcluster of cluster 5.

**Supplementary Table 6. Differentially expressed genes in COVID-19 vs. control sections.** Differentially expressed (DE) genes for COVID-19 versus control lung tissue samples.

**Supplementary Table 7. Differentially expressed genes in COVID-19 vs. control per cluster**. DE genes per cluster for COVID-19 versus control lung tissue samples.

**Supplementary Table 8. Cluster 4 subclustered differentially expressed genes.** Differentially expressed (DE) genes for each subcluster of cluster 4.

**Supplementary Table 9. Cluster 4 subclustered DE genes for COVID-19 vs. control.** DE genes for COVID-19 versus control lung tissue samples for each subcluster of cluster 4.

**Supplementary Table 10. Colocalization analysis results of COVID-19 sections.** Differentially expressed genes in SARS-CoV-2^+^ vs. SARS-CoV-2^-^ spots in COVID-19 sections.

**Supplementary Table 11. SARS-CoV-2 ST gene probe sequences.** Probe sequence information for the ST SARS-CoV-2 gene probes used in the study.
